# Global drivers of diversification in a marine species complex

**DOI:** 10.1101/2019.12.13.874883

**Authors:** Catarina N.S. Silva, Nicholas P. Murphy, James J. Bell, Bridget S. Green, Guy Duhamel, Andrew C. Cockcroft, Cristián E. Hernández, Jan M. Strugnell

## Abstract

Investigating historical gene flow in species complexes can indicate how environmental and reproductive barriers shape genome divergence before speciation. The processes influencing species diversification under environmental change remain one of the central focal points of evolutionary biology, particularly for marine organisms with high dispersal potential. We investigated genome-wide divergence, introgression patterns and inferred demographic history between species pairs of all extant rock lobster species (*Jasus* spp.), a complex with long larval duration, that has populated continental shelf and seamount habitats around the globe at approximately 40°S. Genetic differentiation patterns revealed the effects of the environment and geographic isolation. Species associated with the same habitat structure (either continental shelf or seamount/island) shared a common ancestry, even though the habitats were not adjacent. Differences in benthic temperature explained a significant proportion (41.3%) of the genetic differentiation. The Eastern Pacific species pair of *J. caveorum* and *J. frontalis* retained a signal of strict isolation following ancient migration, whereas species pairs from Australia and Africa and seamounts in the Indian and Atlantic oceans included events of introgression after secondary contact. Parameters estimated for time in isolation and gene flow were congruent with genetic differentiation metrics suggesting that the observed differentiation patterns are the product of migration and genetic drift. Our results reveal important effects of habitat and demographic processes on the divergence of species within the genus *Jasus* providing the first empirical study of genome-wide drivers of diversification that incorporates all extant species in a marine genus with long pelagic larval duration.

## Introduction

The discrete categorization of speciation modes as sympatric, allopatric or parapatric is now considered to be overly simplistic (Butlin et al. 2008). Several events (or modes of speciation) can influence the biogeographic states of populations at different time periods during divergence, and as a result, the speciation process is now generally considered to be gradual and reticulated (Smadja & Butlin 2011; Feder *et al*. 2012). However, the processes responsible for influencing species diversification are still poorly understood and remain one of the central focal points of ecology and evolutionary biology (Arendt *et al*. 2016).

Reconstructing the diversification history for species complexes can be challenging for non-model marine organisms (e.g. Palero *et al*. 2009; Momigliano *et al*. 2017) as many have large population sizes and the potential for long distance dispersal, which frequently result in weak genetic differentiation (Ovenden 2013). For these species, it is often difficult to determine whether weak genetic differentiation is actually present or masked by the large population sizes (Lowe & Allendorf 2010). In addition, marine species with long distance dispersal can quickly fill available niches, leaving fewer opportunities for *in situ* cladogenesis (Pinheiro *et al*. 2017). As a result, only a few studies have reconstructed the history of diversification in marine species (e.g. Crow *et al*. 2010; Le Moan *et al*. 2016; Momigliano *et al*. 2017; Souissi *et al*. 2018; Titus *et al*. 2019).

Changes in the distribution of marine species resulting from historical climatic variation have been an important driver of diversification across taxa (Davis *et al*. 2016). Climatic fluctuations during the late Pleistocene, in particular, resulted in periods of isolation intercalated by contact and gene flow between lineages (Hewitt 2000). These events dramatically transformed available habitat causing major shifts in species distribution ranges and shaping the genetic structure of many marine species worldwide (Benardine Jeffrey *et al*. 2007; Kenchington *et al*. 2009; Van Oppen *et al*. 2011; Strugnell *et al*. 2012). Sequential glacial and interglacial periods have then further shaped the divergence history of species as a result of periods of isolation intercalated by gene flow (Weigelt *et al*. 2016). A better understanding of the species-specific historical context of divergence is therefore needed to estimate the actual timing and role of gene flow during speciation. Understanding how historical climatic fluctuations shaped species divergence provides clues on how species might respond to future environmental changes, which is vital for effective conservation and management plans (Olivieri *et al*. 2016).

Advances in next-generation sequencing (NGS) now provide the opportunity to investigate genome-wide patterns of differentiation along the speciation continuum, allowing the better detection of changes as two lineages diverge from one another on the path to reproductive isolation (Feder *et al*. 2012). In particular, these methods provide effective tools solutions for species with no reference genomes (Catchen *et al*. 2017), which is the case for many marine species including rock lobsters. This technology has also allowed the integration of genomic and environmental data which can be used for testing the hypothesis that selection is more efficient than drift in opposing the homogenizing effects of migration (Manel & Holderegger 2013). In addition, this approach can also detect candidate markers underlying adaptation to local environments for species with moderate to long distance dispersal potential (e.g. Benestan *et al*. 2016; Sandoval-Castillo *et al*. 2018). This robust approach is particularly useful in the marine environment where isolation and speciation is increasingly found to be associated with selection/local adaptation (Rocha *et al*. 2005; Momigliano *et al*. 2017). Improvements in methodology have further enabled the use of genome-wide polymorphism data to infer complex demographic histories and the relative influence of gene flow and historical processes on the genomic landscape. For example, in the marine environment this approach has been used in the European anchovy *Engraulis encrasicolus* (Le Moan *et al*. 2016), the Atlantic Salmon *Salmo salar* (Rougemont & Bernatchez 2018) and the corkscrew sea anemone *Bartholomea annulate* (Titus *et al*. 2019). One increasingly popular approach is demographic inference based on the computation of a joint allele frequency spectrum (JAFS) from genetic polymorphism data (Gutenkunst *et al*. 2009; Excoffier *et al*. 2013). This approach allows an estimation of several demographic parameters such as population sizes, migration rates and time intervals since specific events using a composite likelihood. Therefore, the role of historical events in the diversification and speciation of marine species can now be more accurately determined.

Rock lobsters (*Jasus* spp.) are a useful model to study the role of historical climatic variations and gene flow on divergence. The six extant *Jasus* lobster species (*J. caveorum, J. edwardsii, J. frontalis, J. lalandii, J. paulensis* and *J. tristani*) are distributed in a narrow latitudinal band (∼25° to 47°; Fig. 1) in the Southern Hemisphere (Booth 2006) up to 200 m (Holthuis 1991) and possibly up to 600 m depth (Duhamel personal communication). These animals have a long pelagic larval duration (PLD; up to two years for *J. edwardsii*), with the potential for extensive dispersal (Bradford *et al*. 2015). Despite such a long PLD, all species have a restricted latitudinal distribution; for example, *J. caveorum* is only known from a single seamount in the eastern South Pacific Ocean (Webber & Booth 1995). Phylogenetic relationships between *Jasus* species have been investigated with a limited number of mtDNA markers (Brasher *et al*. 1992; Ovenden *et al*. 1997). Ovenden *et al*. (1997) identified a clade containing *J. edwardsii, J. lalandii* and *J. frontalis*, however, the relative branching order was not resolved by analysis of sequence variation in the cytochrome c oxidase subunit I (COI) and the 16S ribosomal RNA sequences. In addition, the species *J. tristani* and *J. paulensis*, which occur in islands and seamounts off the southern Atlantic and Indian Oceans, respectively, were hypothesized to have come into secondary contact during past glacial periods, resulting in low levels of mtDNA differentiation (Ovenden *et al*. 1997; Groeneveld *et al*. 2012). At the species level, population genetic studies have demonstrated a general pattern of low, yet often significant, differentiation (Ovenden *et al*. 1992; Matthee *et al*. 2007; Porobić *et al*. 2013; Thomas & Bell 2013; Villacorta-Rath *et al*. 2016). Post-settlement selection and chaotic genetic patchiness, also described as a shifting, ephemeral genetic pattern, has also been observed in *J. edwardsii*, highlighting the uncertainties in predicting connectivity between populations of highly dispersive marine organisms (Villacorta-Rath *et al*. 2018).

**Fig. 1.**
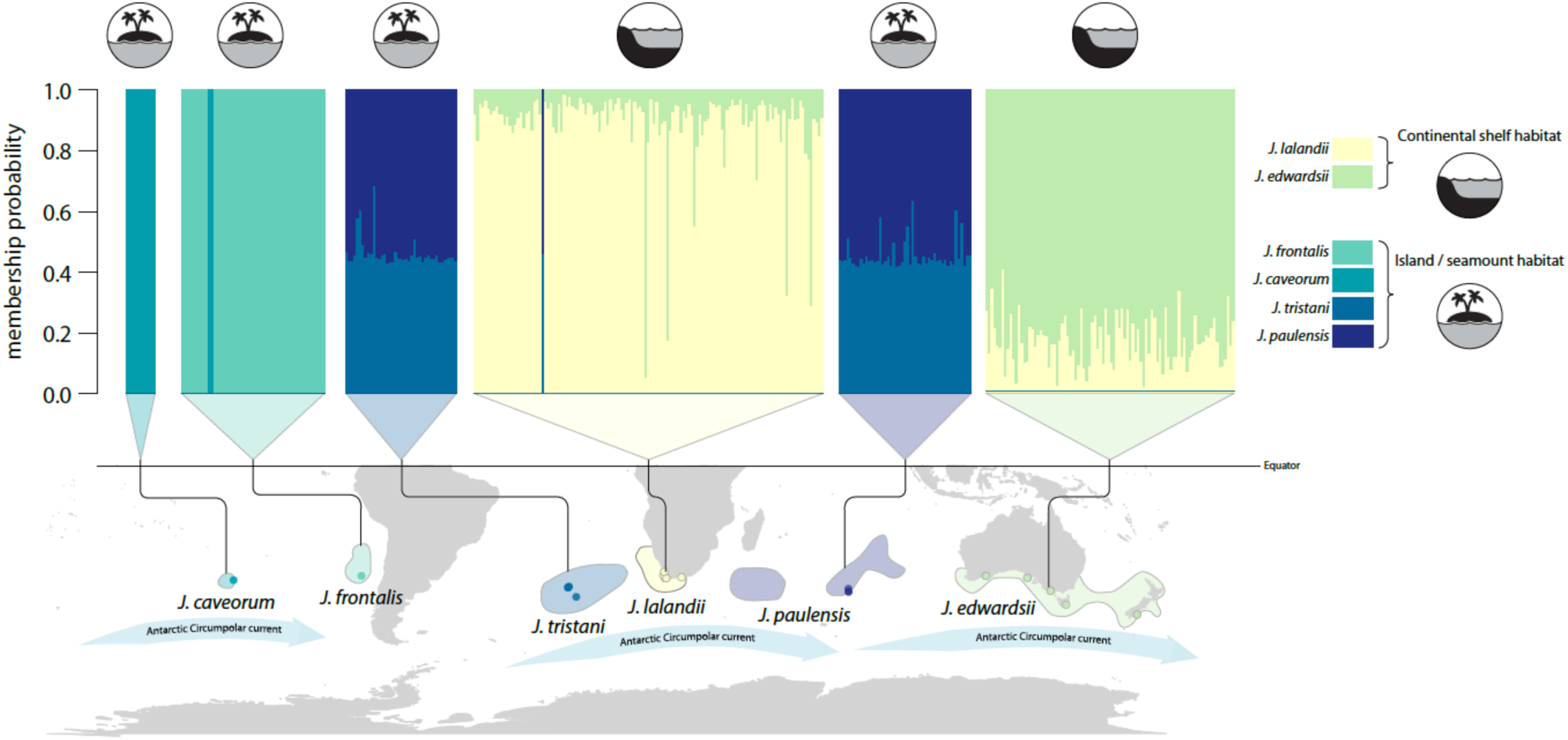
Sample locations, approximate distribution range of *Jasus* spp. (adapted from Booth, 2006) and membership probability plot using 2 principal components.

Although a few studies suggest a recent divergence between *Jasus* lineages (Pollock 1990; Ovenden *et al*. 1997), relatively little attention has focused on investigating diversification processes in *Jasus* lobsters. Here we investigate speciation processes in all the extant lobster species of the genus *Jasus*. This study aims to test for evidence of admixture/introgression between species, investigating the genetic patterns associated with habitat structure (continental shelf or seamount/island) and inferring the demographic history of *Jasus* spp. using genome-wide single nucleotide polymorphisms (SNP).

## Methods

### Sampling, DNA extractions and sequencing

Tissue samples of *Jasus* spp. were collected between 1995 and 2017 from 17 locations throughout the entire genus’ range (Fig. 1). A total of 375 samples were collected (11 *Jasus caveorum*, 53 *J. frontalis*, 41 *J. tristani*, 129 *J. lalandii*, 49 *J. paulensis* and 92 *J. edwardsii*). Tissue was stored in 70% ethanol before processing. Total genomic DNA of *J. caveorum* museum samples was extracted using the QIAamp DNA Micro Kit (Qiagen) according to the manufacturer’s instruction. The remaining tissue samples were extracted using NucleoMag® Tissue (Macherey-Nagel) following to the manufacturer’s instructions.

Library preparation and sequencing was conducted by Diversity Arrays Technology, Canberra, Australia and followed standard protocols of DArTseq™ genotyping technology (Kilian *et al*. 2012). Briefly, approximately 100 ng (2 µL) of each sample was digested with the restriction enzymes PtsI and SphI, and unique barcode sequences simultaneously ligated onto the ends of each resulting fragment as per Kilian et al. 2012. The PstI-compatible adapter included an Illumina flow-cell attachment sequence, a primer sequence and unique barcode, with the reverse SphI-compatible adaptor contained in the flow-cell attachment region. A minimum of 15% random technical replicates were included for downstream quality control. Each sample with fragments containing both PstI and SphI cut sites was amplified in PCR reactions using the following conditions: 94°C for 1 min then 30 cycles of 94 °C for 20 s, 58 °C for 30 s, 72 °C for 45 s, and 72 °C for 7 min. Samples were checked visually on an agarose gel to ensure complete digestion and uniform range of fragment sizes. Using approximately 10 µL of each sample, samples were sequenced on a single flow-cell lane on the Illumina HiSeq2500 for 77 cycles.

### De novo *assembly and variant calling*

Libraries were demultiplexed and reads were filtered for overall quality (–c, –q and –r options) using *process_radtags* in STACKS v.2.0b9 (Catchen *et al*. 2013). The Stacks pipeline *denovo_map*.*pl* was executed to run each of the Stacks modules individually (*ustacks, cstacks, sstacks* and *populations*). The formation of loci was allowed with a maximum of two nucleotides between stacks (M = 2) and a minimum stack depth of three (m = 3) among reads for accurate calling (*ustacks* module). Reads were aligned *de novo* with each other, and a catalogue of putative RAD tags was created (*cstacks* module). Putative loci were searched against the catalog (*sstacks* module) and further filtering was then conducted in the *populations* module.

Retained reads were present in at least 70% of samples within each species, were detected in all species, had a rare allele frequency of at least 2% and had no more than 2 alleles detected. Potential paralogs were excluded by removing markers with heterozygosity > 0.50 within samples and analyses were restricted to one random SNP per locus (using *--write_random_snp*). These filtering steps aimed to exclude as many SNPs as was possible with genotyping errors and missing data.

### Genetic diversity and population structure

Allelic richness, pairwise F_ST_ values and respective p-values were estimated using *hierfstat* package in R (Goudet 2005). The R package *adegenet* was used to estimate observed and expected heterozygosity, inbreeding coefficients and for discriminant analyses of principal components (DAPC) and membership probability plots (Jombart 2008). Outlier analyses were conducted in BayeScan to look for signatures of selection. Prior odds were set to 100 to minimize chances of false positives with 5,000 pilot runs, followed by 100,000 iterations (5,000 samples, a thinning interval of 10, and a burn-in of 50,000).

### Environmental data collection and analyses

Initially, 13 environmental variables were obtained from Bio-Oracle (Assis *et al*. 2018; Table S1). Only uncorrelated variables (r<0.6) were retained resulting in seven layers (surface and benthic temperature mean, surface salinity, surface and benthic current velocity, benthic iron and surface phytoplankton). Multiple regression of distance matrices (MRDM; Legendre et al., 2014) was used to examine the association of geographic distance (estimated as the shortest path distance in the ocean) with patterns of genetic differentiation (measured as pairwise F_ST_ values), using the R package ecodist (Goslee & Urban 2007). Redundancy analysis (RDA; Forester *et al*. 2018) was used to investigate genotype-environment associations using the R package vegan (Oksanen *et al*. 2019). Significance was assessed using a permutation test (999 permutations) for redundancy analysis using the function *anova*.*cca()*.

### Relationships among lineages

The program TREEMIX v1.12 (Pickrell & Pritchard 2012) was used to further investigate historical relationships among lineages. A maximum-likelihood (ML) phylogeny was first inferred and then single migration events between branches were sequentially added to determine whether migration/admixture events improve the likelihood fit. To formally test for admixture between *Jasus* spp., the three-population test (Reich *et al*. 2009) included with TREEMIX was used. In this test, the *f*_*3*_ (X; A,B) statistic is negative when a population X does not form a simple tree with populations A and B, but rather may be a mixture of A and B.

### Demographic modelling

Previous analysis suggests evidence of admixture between species pairs, and so we tested several hypothesis of divergence modes, aiming to identify speciation events through time, for each closely related pair of species: *J. caveorum* - *J. frontalis, J. edwardsii* - *J. lalandii* and *J. tristani* - *J. paulensis*. The species pairs were selected based on their genetic and morphological relationships (Holthuis & Sivertsen 1967; George & Kensler 1970; Brasher *et al*. 1992; Ovenden *et al*. 1997; Groeneveld *et al*. 2012; this study). For each pair, six models were built representing alternative modes of divergence considering possible scenarios: (SI) Strict Isolation, where the environment (e.g. sea level change and ocean currents) promoted allopatry; (IM) Isolation-with-Migration, with continuous gene flow through the speciation process; (AM) Ancient Migration, with an ancient gene flow event but recent isolation; (SC) Secondary Contact, with a recent gene flow event after past isolation; (PAM) Ancient Migration with two periods of gene flow, with two ancient gene flow events but recent isolation; and (PSC) Secondary Contact with two periods of contact, two recent gene flow events after past isolation. Six additional models were built to include expansion/contraction events in the initial models (suffix ‘ex’). All models were implemented allowing for asymmetric migration rates (m12, m21).

Demographic inference was performed using the diffusion approximation method implemented in the software ∂a∂i (Gutenkunst *et al*. 2009). The function *vcf2dadi* in the R package radiator (Gosselin 2017) was used to create ∂a∂i SNP input files. We used the folded joint site frequency spectrum (JSFS) for model selection because the closest out-group (*Sagmariasus verreauxi*) was too distant (diverged around 11 Mya; Ovenden *et al*. 1997), which resulted in a highly reduced number of orientable polymorphisms.

In total, 12 models were tested per species pair, fitted with the observed joint site frequency spectrum (SFS) using 20 replicate runs per model and the best model was retained (Fig. S4). The Akaike information criterion (AIC) was used to perform comparisons among models (Sakamoto *et al*. 1986).

To compare among nested models of increasing complexity and address over-parametrization issues we used the comparative framework of Tine *et al*. (2014) by penalizing models which contain more parameters. For each species pair, a score was estimated for each model using:

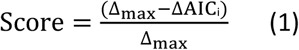

where, Δ_max_ corresponds to the difference in AIC between the worst and the best performing model (Δ_max_ = AIC_max_ - AIC_min_) and ΔAIC_i_ = AIC_i_ - AIC_min_. Therefore, the worst model has a score of 0 and the best model has a score of 1. To evaluate the relative probabilities of the different models within each species pair, Akaike weights (W_AIC_) were also calculated following:

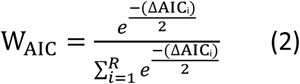

where R corresponds to the total number of models considered (R=12).

Demographic parameters were converted into indicative biologically units, given the missing crucial information about mutation rate per generation in *Jasus* spp. The ancestral effective population size (N_ref_) before split for each species pair was estimated following:

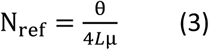

with θ being the optimal multiplicative scaling factor, μ the mutation rate (fixed at 8×10^−8^ mutations per site per generation; Obbard *et al*. 2012) and L the effective length of the genome explored:

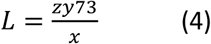

where x is the number of SNPs originally detected from y RAD-tags of 73 bp present in the initial data set, and z the number of SNPs retained, following Rougeux *et al*. (2017). Estimated units in 2N_ref_ were converted to years assuming a generation time of 10 years (Pecl *et al*. 2009). Estimated migration rates were divided by 2N_ref_ to obtain the proportion of migrants in every generation.

## Results

### Genetic diversity and population structure

Sequencing yielded a total of 1,501,921,855 quality-trimmed sequencing reads, providing an average depth of coverage per individual over all SNPs of 58.9x. After applying the different filtering steps, 2,596 SNPs common to all species were retained for subsequent analyses. The lowest values of observed heterozygosity, expected heterozygosity, and allelic richness were observed for *J. caveorum*. While *J. frontalis* had the highest inbreeding coefficients (0.502), all species had very similar values. The highest values of allelic richness were observed for *J. lalandii* (Table 1). The highest pairwise F_ST_ values were observed for *J. tristani* – *J. caveorum* and *J. paulensis* – *J. caveorum* (F_ST_ = 0.463 and F_ST_ =0.436, respectively, p < 0.05), while the lowest values were observed for *J. tristani* – *J. paulensis* (F_ST_ = 0.022, p < 0.01; Table 2).

**Table 1.**
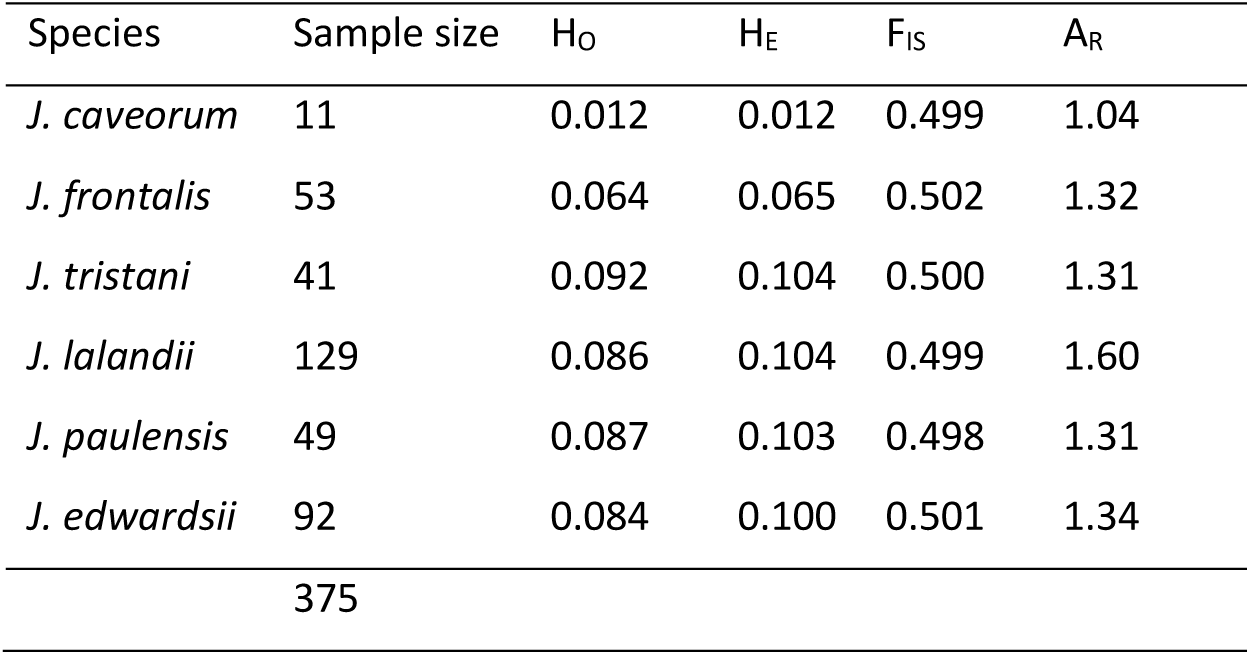
Summary statistics of genetic diversity per species using 2,596 SNPs. H_O_: observed heterozygosity, H_E_: expected heterozygosity, F_IS_: inbreeding coefficient, A_R_: allelic richness

**Table 2.** Pairwise F_ST_ values (below diagonal) and corresponding p-values (above diagonal) estimated using *hierfstat* package in R.

|  | *J. caveorum* | *J. frontalis* | *J. tristani* | *J. lalandii* | *J. paulensis* | *J. edwardsii* |
| --- | --- | --- | --- | --- | --- | --- |
| *J. caveorum* | 0 | 0.010 | 0.013 | 0.010 | 0.011 | 0.012 |
| *J. frontalis* | 0.081 | 0 | 0.007 | 0.005 | 0.006 | 0.007 |
| *J. tristani* | 0.463 | 0.206 | 0 | 0.007 | 0.008 | 0.009 |
| *J. lalandii* | 0.305 | 0.137 | 0.387 | 0 | 0.006 | 0.007 |
| *J. paulensis* | 0.436 | 0.229 | 0.022 | 0.408 | 0 | 0.007 |
| *J. edwardsii* | 0.413 | 0.106 | 0.441 | 0.230 | 0.452 | 0 |

No signatures of selection were detected by the outlier detection analyses (Fig. S1). Lobster species were grouped into three main clusters by discriminant analyses of principal components when using 2 PCs (52.3% variation) (Fig. 2). There was evidence of admixture, in particular between *J. paulensis* - *J. tristani* in the membership probability plot and the DAPC results, and pairwise F_ST_ values (Fig. 1 and 2). The first DAPC axis (LD1) explained 29.9% of the variation and highlighted the divergence between habitat structure (i.e. *J. edwardsii* and *J. lalandii* vs. remaining species; Fig. S2a), while the second DAPC axis (LD2), which explained 22.4% of the variation, showed three main clusters and highlighted the differences between *J. paulensis* and *J. tristani* and the remaining species (Fig. S2b).

**Fig. 2.**
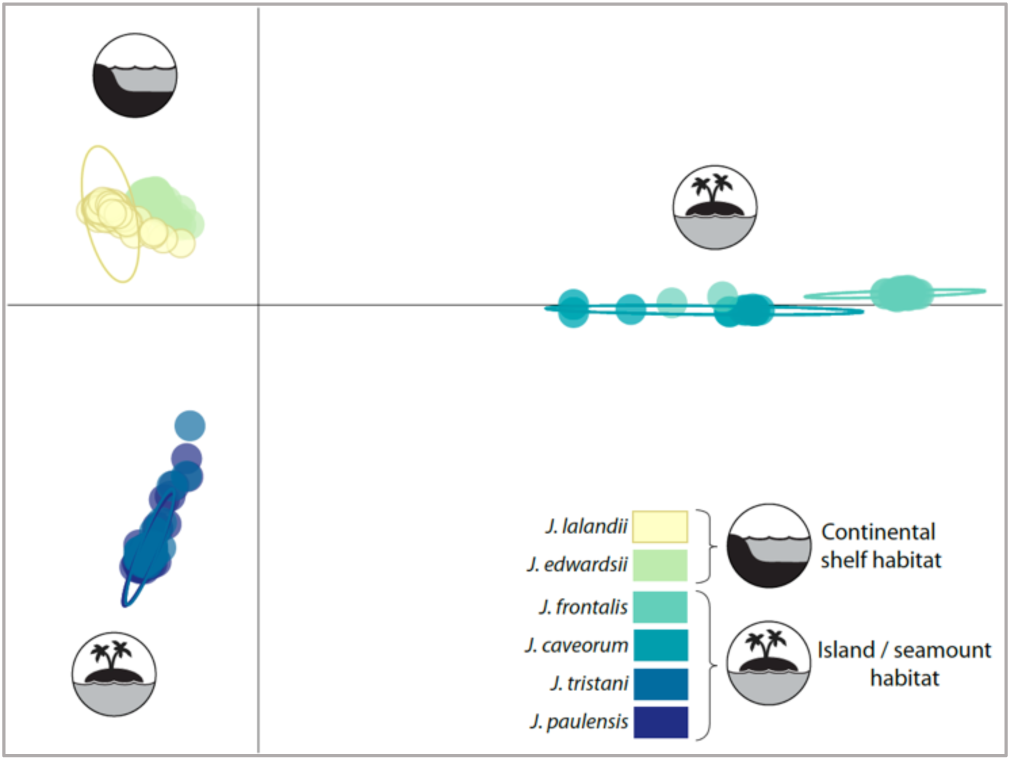
Discriminant analyses of principal components (DAPC) of *Jasus* spp. using 2 principal components (explaining 52.3% variation).

### Genotype-environment associations

Geographic distance explained 23.3% of the total genetic variation (F_ST_) between rock lobster species (p<0.01), using multiple regression of distance matrices (MRDM). The model with all seven environmental variables explained 52.9% of the total genetic variation between lobster species (p<0.001), while the model with the four most important environmental variables explained 51.4% of the total genetic variation (p<0.001). Benthic mean temperature was the single environmental variable that explained most of the genetic differentiation while controlling for the effects of geographic distance (41.3%; p<0.001; Table 3).

**Table 3.** Summary of the models for the multiple regression of distance matrices (MRDM) analyses using F_ST_ as a measure of genetic differentiation. GeoDist: geographic distance (km); SurfTemp and BenTemp: Surface and benthic temperature (°C); SurfSal: Surface salinity (PSS); SurfCurren and BenCurren: Surface and benthic current velocity (m.s^-1^); BenIron: Benthic dissolved iron (mmol.m^-3^); SurfPhyto: surface phytoplankton (mmol.m^-3^).

| Factors | $r^2$ | p-value |
| --- | --- | --- |
| GeoDist | 0.233 | <0.01 |
| GeoDist + SurfTemp | 0.235 | <0.001 |
| GeoDist + BenTemp | 0.413 | <0.001 |
| GeoDist + SurfSal | 0.233 | <0.001 |
| GeoDist + SurfCurren | 0.238 | <0.01 |
| GeoDist + BenCurren | 0.298 | <0.001 |
| GeoDist + BenIron | 0.237 | <0.001 |
| GeoDist + SurfPhyto | 0.290 | <0.001 |
| GeoDist + SurfTemp + BenTemp + SurfSal + SurfCurren + BenCurren + BenIron + SurfPhyto | 0.529 | <0.001 |
| GeoDist + SurfTemp + BenTemp + BenCurren + SurfPhyto | 0.514 | <0.001 |

All seven environmental variables explained 18% of the variation in rock lobster species (p<0.001) when using the constrained ordination in RDA analyses. All values of the variance inflation factors were below five, indicating that multicollinearity among the predictor variables is not inflating the model. The first five constrained axes were significant in explaining the genetic variation between species (each explaining 53.3%, 25.3%, 12%, 5.2% and 2.2%, respectively; p<0.001; Fig. 3). Genetic variation of *J. caveorum* and *J. frontalis* was associated with higher surface temperature, while *J. paulensis* and *J. tristani* were associated with lower surface temperature. *J. edwardsii* and *J. lalandii* were associated with higher benthic temperature, benthic current velocity and benthic iron. Finally, *J. lalandii* was associated with higher surface phytoplankton, while *J. edwardsii* was associated with higher surface current velocity (Fig. 3).

**Fig. 3.**
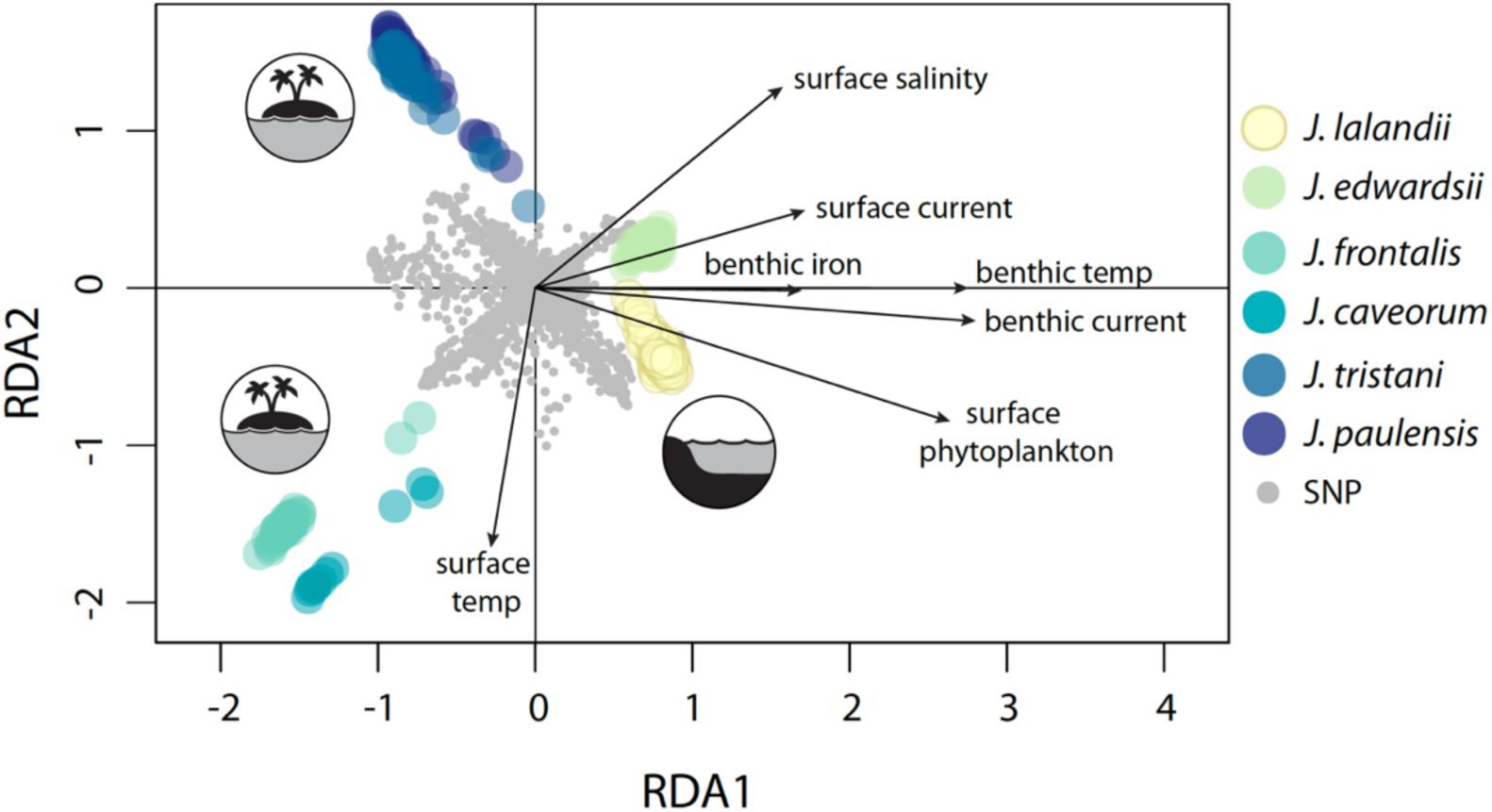
Ordination plot of redundancy analysis (RDA) of *Jasus* spp. The vectors are the environmental predictors (see Table 3 for a detailed description).

### Relationships among lineages

Results from TREEMIX identified three ancestral events of admixture (Fig. S3). However, from the three-population test of admixture, only two *f*_*3*_ values were negative with associated Z-scores < −0.6, indicating evidence that *J. tristani* does not form a simple tree with *J. paulensis, J. lalandii* and *J. edwardsii*, but rather may be a mixture of these (Table S2, Supporting information). Therefore, the three-population test supported the ancestral event of admixture detected by TREEMIX from the most recent common ancestor (MRCA) of *J. lalandii* and *J. edwardsii* to *J. tristani*. The genetic relationships among species inferred by TREEMIX revealed similar patterns to the genetic differentiation analyses, clearly separating species pairs *J. lalandii* – *J. edwardsii, J. paulensis* – *J. tristani* and *J. caveorum* – *J. frontalis* (Fig. S3).

### Demographic modelling

In general, SC and PSC models provided better fits to the data with good predictions of the joint site frequency spectrum (JSFS) asymmetry for the *J. paulensis* – *J. tristani* and *J. edwardsii* – *J. lalandii* pairs while AM, PAM and SI had better support for the *J. caveorum* – *J. frontalis* pair (Table S3, Fig. S4). Incorporating population expansion events (suffix ‘ex’) improved the fit of PSC models for all species pairs but there was not a clear pattern for PAM models. In contrast, the strict isolation (SI) and ancient migration (AM, PAM) models were weakly supported for the *J. paulensis* – *J. tristani* and *J. edwardsii* – *J. lalandii* pairs while the secondary contact models (SC, PSC) were weakly supported for the *J. caveorum* – *J. frontalis* pair (Table S3, Fig. S4).

Asymmetries in gene flow with ratios of m_21_/m_12_ indicated a stronger migration from population two to population one in all species pairs, and the lower proportion of migrants was observed for the *J. edwardsii – J. lalandii* pair (Fig. 4, Table 4). Detailed results for demographic inferences are provided in Table S3, Fig. 4, Table 4, Fig. S4 and Fig. S5 (supporting information).

**Table 4.** Parameters estimates for the best model of each species pair with standard deviation. *J. paulensis* – *J. tristani*: secondary contact with two periods of contact and recent population expansion (PSCex); *J. edwardsii – J. lalandii*: secondary contact and recent population expansion (SCex); *J. caveorum* – *J. frontalis*: ancient migration with two periods of ancient gene flow and recent population contraction (PAMex).

| Species group | 1: *J. paulensis*,<br>2: *J. tristani* | 1: *J. edwardsii*,<br>2: *J. lalandii* | 1: *J. caveorum*,<br>2: *J. frontalis* |
| --- | --- | --- | --- |
| **Best Model** | PSCex | PSCex | PAMex |
| **K** | 9 | 9 | 9 |
| **N<sub>ref</sub>** | 43.96 | 85.64 | 64.19 |
| **N<sub>1</sub>** | 3.50 ± 0.84 | 1.50 ± 0.25 | 1.30 ± 0.35 |
| **N<sub>2</sub>** | 49.67 ± 19.38 | 41.01 ± 15.61 | 19.19 ± 9.09 |
| **N<sub>1e</sub>** | 0.67 ± 0.16 | 2.44 ± 0.41 | 0.05 ± 0.24 |
| **N<sub>2e</sub>** | 2.52 ± 0.53 | 1.28 ± 0.34 | 0.20 ± 0.23 |
| **m<sub>12</sub>** | 11.00 ± 3.77 | 1.77 ± 0.23 | 7.71 ± 2.33 |
| **m<sub>21</sub>** | 2.59 ± 2.22 | 0.09 ± 0.16 | 1.99 ± 0.36 |
| **T<sub>S</sub>** | 11.94 ± 2.67 | 10.66 ± 3.37 | 9.74 ± 4.48 |
| **T<sub>SC/AM</sub>** | 2.20 ± 0.70 | 0.27 ± 0.10 | 1.20 ± 0.19 |
| **T<sub>e</sub>** | 0.51 ± 0.20 | 0.29 ± 0.11 | 0.24 ± 0.27 |
| **T<sub>total</sub>** | 29.29 ± 7.15 | 22.46 ± 7.15 | 22.36 ± 9.87 |
| **\*m<sub>12</sub>** | 0.12 ± 0.04 | 0.01 ± 0.001 | 0.06 ± 0.02 |
| **\*m<sub>21</sub>** | 0.03 ± 0.03 | 0.00 ± 0.00 | 0.01 ± 0.00 |
| **\*T<sub>S</sub>** | 10,524 ± 2,346 | 18,258 ± 5,769 | 12,507 ± 5,746 |
| **\*T<sub>SC/AM</sub>** | 1,941 ± 618 | 470 ± 167 | 1,541 ± 239 |
| **\*T<sub>e</sub>** | 447 ± 179 | 503 ± 185 | 306 ± 352 |
| **\*T<sub>total</sub>** | 25,826 ± 6,286 | 38,460 ± 12,242 | 28,709 ± 12,674 |

K: The number of free parameters in the model
N<sub>ref</sub>: The effective size of the ancestral population before the split
N<sub>1</sub>: The effective size of population 1 before expansion
N<sub>2</sub>: The effective size of population 2 before expansion
N<sub>1e</sub>: The effective size of population 1 after expansion
N<sub>2e</sub>: The effective size of population 2 after expansion
m<sub>12</sub>: The neutral movement of genes from population 2 to population 1 in units of 2N<sub>ref</sub> generations
m<sub>21</sub>: The neutral movement of genes from population 1 to population 2 in units of 2N<sub>ref</sub> generations
T<sub>S</sub>: The time of divergence in strict isolation in units of 2N<sub>ref</sub> generations

**Fig. 4.**
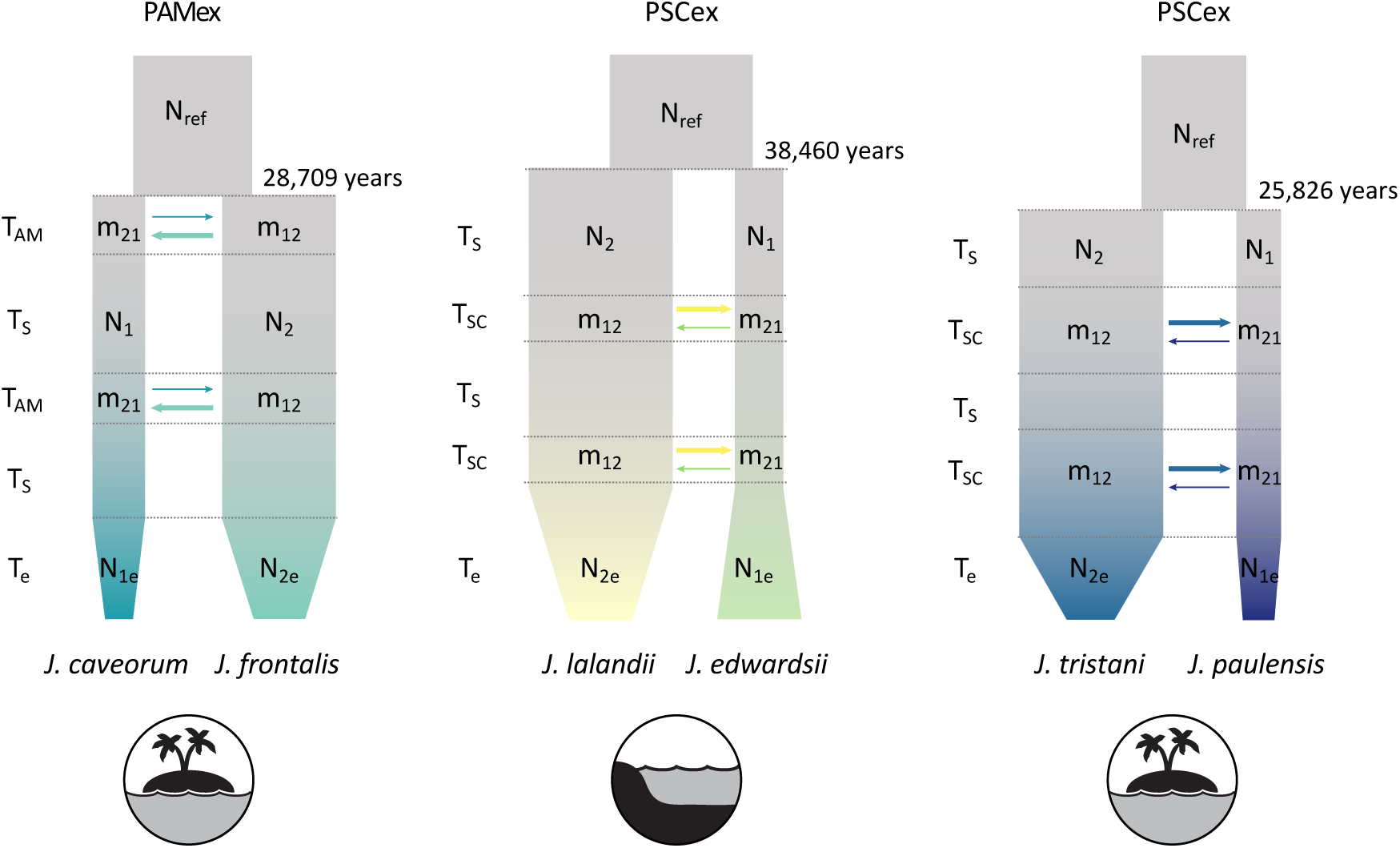
Representation of the best demographic model for each species pair; *J. caveorum* – *J. frontalis*: ancient migration with two periods of ancient gene flow and recent population contraction (PAMex); *J. lalandii* – *J. edwardsii* and *J. tristani* – *J. paulensis*: secondary contact with two periods of contact and recent population expansion (PSCex). Asymmetric migration rates (m_21_ and m_12_) are represented by the arrows with higher rates of migration from population two to population one for all species pairs (thicker lines in arrows). Width of the boxes represent sizes of the ancestral population (N_ref_), population sizes before expansion/contraction (N_1_, N_2_) and population sizes after expansion/contraction (N_1e_, N_2e_). T_s_ is the time of divergence in strict isolation, T_SC/AM_ the time of secondary contact or ancient migration and T_e_ the time of expansion.

The best supported model for the *J. paulensis* – *J. tristani* pair was PSCex (Table S3). Within this model, total divergence time between species was approximately 25,826 ± 6,286 years ago (Table 4). The period without contact was approximately 5.4 times longer than the period with secondary contact. The best supported model for the *J. edwardsii* – *J. lalandii* pair was PSCex (Table S3). Total divergence time between *J. edwardsii* and *J. lalandii* was approximately 38,460 ± 12,242 years ago (Table 4). The period without contact was approximately 38.8 times longer than the period with secondary contact. Finally, the best supported model for the *J. caveorum* – *J. frontalis* pair was PAMex (Table S3). Within this model, total divergence time between species was approximately 28,709 ± 12,674 years (Table 4) and the period without contact was approximately 8.1 times longer than the period with ancient migration. Therefore, divergence times with errors overlap across the three species pairs and was estimated to be between 19,540 and 32,112 years for *J. paulensis* – *J. tristani*, 26,218 and 50,702 years for *J. edwardsii* – *J. lalandii* and 16,035 and 41,383 years for *J. caveorum* – *J. frontalis*.

## Discussion

Here we investigated genome-wide divergence and introgression patterns in all extant species of rock lobsters (*Jasus* spp.) for the first time. Genetic differentiation patterns revealed the effects of the environment and geographical isolation. Species that were associated with the same habitat structure (continental shelf or seamount/island) were more closely related to each other than with species from a different habitat structure. Benthic temperature was the single environmental variable that explained most of the genetic differentiation (F_ST_) while controlling for the effects of geographic distance (41.3%), and *J. edwardsii* and *J. lalandii* were associated with higher mean benthic temperatures. We also detected multiple introgression events (gene flow) since the first divergence in all species pairs.

### Divergence during Pleistocene

Divergence times between species pairs estimated by demographic modelling overlap across all species comparisons and suggest that global/widespread processes might have driven initial divergence across all species. During the divergence period estimated for *Jasus* spp. (from 38,460 to 25,826 years ago) global temperature and sea levels were decreasing (Clark *et al*. 2009). Sea level was 65 to 125 metres lower than today possibly creating more shallow benthic habitat for lobsters in the open ocean in contrast with a reduction in the available continental shelf habitat (Schaaf 1996). A significant increase in the Southern Ocean barotropic stream function occurred between 49,000 and 28,500 years ago (Fogwill *et al*. 2015), which may have increased the dispersal potential of larvae. Also, a large increase in subantarctic productivity occurred 35–65,000 years ago during Heinrich events H4–H6 (Sachs & Anderson 2005), which may have enabled planktotrophic larvae to survive for longer periods of time. These processes may have facilitated worldwide dispersal and colonization of circumpolar habitats by the *Jasus* spp. ancestor during this period (Fig. 5).

**Fig. 5.**
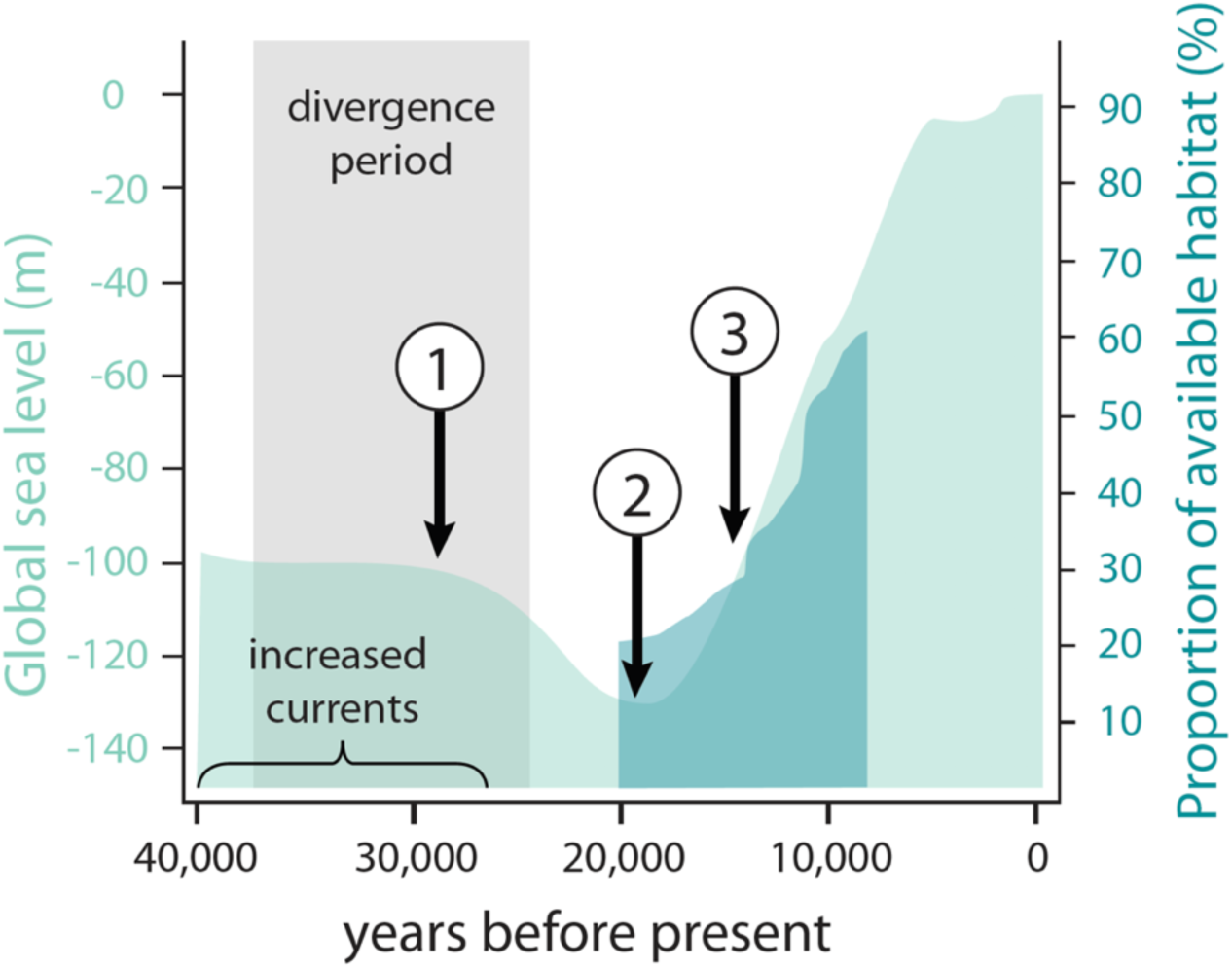
Changes in global sea level (adapted from Huybrechts 2002) and proportion of available habitat in the photic zone (Schaaf 1996) relative to present day. 1) Antarctic sea ice expands around 28,500 years ago (Fogwill *et al*. 2015); 2) Ocean currents reduction (around 22,000 years ago; Alloway *et al*. 2007); 3) West Antarctic Ice Sheet deglaciation (around 14,500 years ago; Clark *et al*. 2009).

During the last glacial maximum (LGM) about 19,000-22,000 years ago, temperature and sea levels reached minimum values (Clark *et al*. 2009). New ecological zones could have appeared, favouring species associated with seamount and island habitats, which could explain the ancient secondary contact events for *J. caveorum* and *J. frontalis* (Schaaf 1996). On the other hand, there was a shrinkage of the shallow continental margins habitat and species therein (Schaaf 1996; Holland 2012). Species associated with continental shelf habitat could also have shifted northwards while tracking their thermal optima. These changes in habitat may have resulted in longer periods of isolation between the continental shelf associated species *J. lalandii* and *J. edwardsii*.

Transition from the LGM to the Holocene precipitated further changes in the available shallow benthic habitat for lobsters with the increase in sea levels and temperature (Clark *et al*. 2009). There were two major expansions of available shallow benthic habitat at 14,000 and 11,500 years (Schaaf 1996) which, for example, could have increased the suitable habitat for *J. frontalis*, having a positive effect on the population size and the effective population size (Porobić *et al*. 2013). However, the variation of the photic sea-bottom area was not linear nor directly correlated with the sea-level oscillations, but reflected topography patterns (Schaaf 1996). These fluctuations in available habitat resulted in alternating periods of isolation and gene flow that have shaped the present genetic signatures of *Jasus* lobsters (Fig. 5).

### Are species genetically isolated?

The present study suggests that global processes might have driven initial divergence across all *Jasus* species. However, current genetic signatures highlight the complexities exclusive to each species evolution. For example, *J. edwardsii* and *J. lalandii* were the most genetically differentiated species pair (F_ST_=0.230), followed by *J. caveorum* – *J. frontalis* (F_ST_=0.081) and *J. tristani* – *J. paulensis* (F_ST_=0.022). This was in agreement with the parameters estimated from demographic modelling, in particular the period of gene flow estimated for each species pair (approximately 470 ± 167 years, 1,541 ± 239 years and 1,941 ± 618 years, respectively).

Our study provides genome-wide evidence of admixture between *J. paulensis* - *J. tristani* that also showed great genetic similarity. Although George & Kensler (1970) have noted that *J. tristani* and *J. paulensis* possess a significant difference in the abdominal sculpture index, genetic evidence suggests that these species can be synonymized as *J. paulensis*, which was also proposed by Groeneveld *et al*. (2012) using the mitochondrial cytochrome oxidase I gene. As a comparison, since initial divergence, *J. tristani* and *J. paulensis* spent 4.1 times longer in secondary contact than *J. edwardsii* – *J. lalandii* and 1.2 times longer than *J. caveorum* and *J. frontalis*. The Tristan da Cunha and Gough Islands (current distribution of *J. tristani*) and the Amsterdam and St. Paul Islands (current distribution of *J. paulensis*) have been grouped in the same zoogeographic province (called the West Wind Drift Islands Province) based on endemic fish fauna distribution (Collette & Parin 1991). Therefore, the distribution of marine species with a pelagic larvae stage may be influenced by the currents of the West Wind Drift and the long periods of gene flow may explain the close relationship between *J. tristani* and *J. paulensis*.

Species associated with continental shelf habitat *J. edwardsii* and *J. lalandii* were genetically more closely related to each other than to *J. tristani* and *J. paulensis* (island/seamount habitat) despite their geographic locations (i.e. *J. lalandii* is geographically closer to *J. tristani* and *J. paulensis* than to *J. edwardsii*; Fig. 1). Although connectivity would be possible between these species groups given the dispersal potential (indeed, *J. lalandii* larvae have been found in the southwest Indian Ocean as far east as Amsterdam Island, adjacent to the *J. paulensis* habitat (Booth & Ovenden 2000)), species appear to be adapted to local environmental conditions. Our results show that benthic temperature might be a limiting factor affecting gene flow between species from island/seamount and continental shelf habitat. Temperature is important for regulating the rate of embryological development in lobsters (Phillips 2013) and could limit reproduction of species adapted to local benthic temperatures.

It has been shown that during the post-larval or puerulus stage, *J. edwardsii* are able to recognize environmental cues such as chemical, acoustic and substrate cues and increase settlement success in suitable habitats (Stanley *et al*. 2015; Hinojosa *et al*. 2016, 2018). Magnetism is also an important cue for adult Western Rock Lobster, *Panulirus cygnus* (Lestang 2014) and adults and postlarvae of spiny lobster *Panulirus argus* (Boles & Lohmann 2003; Kough *et al*. 2014; Ernst & Lohmann 2016). Our results show that genetic diversity of *J. lalandii* and *J. edwardsii* is associated with benthic iron. Therefore, it is possible that *Jasus* larvae are able to use environmental cues such as magnetism for orientation and different species have adapted to recognize local ecotypes and settle on habitats with contrasting structure. Ovenden *et al*. (1997) also suggested that the common ancestor of *J. tristani* and *J. paulensis* may have been able to recognize environmental cues from island and seamount habitats that allowed them to colonize mid-ocean habitats during glacial periods.

In highly dispersive marine taxa, interglacial recolonization of high-latitude habitats can occur rapidly (Hewitt 2000). Such patterns have been established for a range of Northern Hemisphere marine species (e.g. Marko 2004; Ledoux *et al*. 2018), but relatively little is known about the genetic effects of recent glaciations in the Southern Hemisphere (but see e.g. Fraser *et al*. 2009; Strugnell *et al*. 2012; Porobić *et al*. 2013). This study revealed genome-wide patterns of divergence and introgression in all extant species of a highly dispersive marine taxa for the first time. While results point to the important role of demographic and neutral processes of differentiation between species pairs, it also suggests a possible effect of selection in promoting genetic divergence between habitats. Future studies should address the role of adaptive processes to elucidate their relative contribution in shaping genome divergence and speciation of *Jasus* lobsters and to better understand how future environmental change will impact species distribution.

## Data availability

Raw demultiplexed sequencing data will be available at Dryad. Pipelines for *de novo* assembly, genetic structure, environmental association and demographic inference analyses will be available at github after publication.

## Acknowledgments

Funding for this research was provided by an Australian Research Council Discovery Project grant (Project No. DP150101491) awarded to J.M.S., J.J.B., B.S.G. and N.P.M. We would like to thank Gary Carlos (University of Tasmania), Colin Fry (University of Tasmania), Daniel Ierodiaconou (Deakin University), Kent Way, Andrew Kent, Geoff Liggins, Marcus Miller, Giles Ballinger, Darrel Sykes (DAFF), Rick Webber (Te Papa Museum), Jason How (Department of Fisheries, Western Australia) and Sadie Mills (NIWA) for field assistance and sample collection; T.A.A.F. (Terres Australes et Antarctiques Françaises), for their French fisheries observer service “COPEC”, the fishery observer Sophie Cascade on board the F.V. “AUSTRAL” and Charlotte Chazeau to have made available biological scientific samples and data of *Jasus paulensis* from catches in the Saint-Paul/Amsterdam French EEZ; the help of crew has also been appreciated. C.E.H was supported by FONDECYT grants #1170815.

## Author contributions

All authors contributed insights about data analysis, interpretation of results and edited the final drafts of the manuscript. C.N.S.S. analysed the data. C.N.S.S., J.M.S and N.P.M. conceived the study.

