## Supporting information for "Global drivers of diversification in a marine species complex"

### Supplementary Information

**Table S1.** Environmental variables (layer names and units) obtained from Bio-Oracle (Assis *et al.* 2018)

| Layer | Units |
|---|---|
| Temperature (surface and benthic) | $^{o}C$ |
| Salinity (surface and benthic) | PSS |
| Current velocity (surface and benthic) | $m.s^{-1}$ |
| Chlorophyll (surface and benthic) | $mg.m^{-3}$ |
| Dissolved iron (surface and benthic) | $mmol.m^{-3}$ |
| Calcite (surface) | $mol.m^{-3}$ |
| Phytoplankton (surface and benthic) | $mmol.m^{-3}$ |

**Table S2.** Results from the three-population test (Reich *et al.* 2009). Significant values are in bold.

| | $f_3$ stat | SE $f_3$ stat | Z-score |
|---|---|---|---|
| **JTR;JLA,JPA** | **-0.0003** | **0.001** | **-0.590** |
| **JTR;JED,JPA** | **-0.0002** | **0.001** | **-0.336** |
| JTR;JFR,JPA | 0.0002 | 0.001 | 0.290 |
| JTR;JCA,JPA | 0.0004 | 0.001 | 0.587 |
| JPA;JCA,JTR | 0.003 | 0.001 | 5.072 |
| JPA;JFR,JTR | 0.003 | 0.001 | 5.578 |
| JPA;JED,JTR | 0.004 | 0.001 | 6.384 |
| JPA;JLA,JTR | 0.004 | 0.001 | 6.846 |
| JLA;JED,JTR | 0.025 | 0.003 | 9.266 |
| JLA;JED,JPA | 0.025 | 0.003 | 9.267 |
| JLA;JED,JFR | 0.032 | 0.003 | 10.775 |
| JED;JCA,JLA | 0.032 | 0.003 | 11.937 |
| JLA;JCA,JED | 0.033 | 0.003 | 10.879 |
| JED;JFR,JLA | 0.033 | 0.003 | 12.176 |
| JED;JLA,JPA | 0.040 | 0.003 | 14.271 |
| JED;JLA,JTR | 0.040 | 0.003 | 14.411 |
| JFR;JCA,JLA | 0.042 | 0.004 | 11.943 |
| JFR;JCA,JED | 0.043 | 0.004 | 12.050 |
| JFR;JCA,JTR | 0.044 | 0.004 | 12.405 |
| JFR;JCA,JPA | 0.044 | 0.004 | 12.425 |
| JCA;JFR,JPA | 0.052 | 0.004 | 13.322 |
| JCA;JFR,JTR | 0.052 | 0.004 | 13.363 |
| JCA;JED,JFR | 0.053 | 0.004 | 13.480 |
| JCA;JFR,JLA | 0.054 | 0.004 | 13.646 |
| JLA;JFR,JTR | 0.070 | 0.004 | 15.645 |
| JLA;JFR,JPA | 0.070 | 0.005 | 15.632 |
| JLA;JCA,JTR | 0.072 | 0.005 | 15.741 |

| | | | |
|---|---|---|---|
| JLA;JCA,JPA | 0.073 | 0.005 | 15.752 |
| JTR;JCA,JLA | 0.077 | 0.005 | 17.118 |
| JED;JFR,JTR | 0.078 | 0.005 | 17.118 |
| JED;JFR,JPA | 0.078 | 0.005 | 17.069 |
| JED;JCA,JTR | 0.079 | 0.005 | 17.044 |
| JTR;JFR,JLA | 0.079 | 0.004 | 17.668 |
| JED;JCA,JPA | 0.080 | 0.005 | 17.023 |
| JPA;JCA,JLA | 0.081 | 0.005 | 17.556 |
| JPA;JFR,JLA | 0.083 | 0.005 | 18.157 |
| JTR;JCA,JED | 0.085 | 0.005 | 17.973 |
| JTR;JED,JFR | 0.086 | 0.005 | 18.306 |
| JPA;JCA,JED | 0.088 | 0.005 | 18.313 |
| JPA;JED,JFR | 0.090 | 0.005 | 18.727 |
| JTR;JED,JLA | 0.124 | 0.005 | 23.336 |
| JPA;JED,JLA | 0.128 | 0.005 | 23.695 |
| JFR;JED,JPA | 0.129 | 0.006 | 22.898 |
| JFR;JED,JTR | 0.129 | 0.006 | 22.979 |
| JFR;JLA,JPA | 0.136 | 0.006 | 23.773 |
| JFR;JLA,JTR | 0.136 | 0.006 | 23.931 |
| JCA;JED,JPA | 0.138 | 0.006 | 22.350 |
| JCA;JED,JTR | 0.139 | 0.006 | 22.417 |
| JCA;JLA,JPA | 0.146 | 0.006 | 23.294 |
| JCA;JLA,JTR | 0.146 | 0.006 | 23.446 |
| JLA;JPA,JTR | 0.150 | 0.006 | 26.307 |
| JED;JPA,JTR | 0.164 | 0.006 | 27.900 |
| JLA;JCA,JFR | 0.165 | 0.006 | 25.709 |
| JED;JCA,JFR | 0.165 | 0.006 | 25.837 |
| JTR;JCA,JFR | 0.172 | 0.007 | 26.299 |
| JFR;JED,JLA | 0.174 | 0.006 | 27.249 |
| JPA;JCA,JFR | 0.175 | 0.007 | 26.486 |
| JCA;JED,JLA | 0.186 | 0.007 | 26.691 |
| JFR;JPA,JTR | 0.216 | 0.007 | 31.292 |
| JCA;JPA,JTR | 0.223 | 0.007 | 30.349 |

**Table S3.** Results of model fitting for models of divergence for each species pair: Strict Isolation (SI), Isolation-with-Migration (IM), Ancient Migration (AM), Secondary Contact (SC), Ancient Migration with two periods of ancient gene flow (PAM) and Secondary Contact with two periods of contact (PSC); and additional six models including population expansion (prefix 'ex').

| Group | Model | K | LogL | AIC | $\Delta AIC_i$ | Score | $W_{AIC}$ | Theta |
|---|---|---|---|---|---|---|---|---|
| JPA-JTR | PSCex | 9 | -678.7 | 1375.5 | 0.0 | 1.00 | 1 | 11.1 |
| | SCex | 9 | -704.8 | 1427.6 | 52.1 | 0.98 | 0 | 55.1 |
| | PSC | 6 | -749.1 | 1510.1 | 134.6 | 0.96 | 0 | 23.3 |
| | SC | 6 | -758.3 | 1528.6 | 153.1 | 0.95 | 0 | 74.5 |
| | SI | 3 | -772.2 | 1550.4 | 174.9 | 0.94 | 0 | 223.5 |
| | IM | 5 | -787.7 | 1585.4 | 209.9 | 0.93 | 0 | 80.1 |
| | IMex | 8 | -841.1 | 1698.2 | 322.7 | 0.89 | 0 | 231.4 |
| | SIex | 6 | -886.9 | 1785.8 | 410.3 | 0.87 | 0 | 251.0 |
| | PAM | 6 | -1146.3 | 2304.6 | 929.1 | 0.70 | 0 | 16.9 |
| | AM | 6 | -1232.3 | 2476.7 | 1101.2 | 0.64 | 0 | 38.1 |
| | PAMex | 9 | -1890.9 | 3799.8 | 2424.3 | 0.21 | 0 | 53.7 |
| | AMex | 9 | -2205.1 | 4428.2 | 3052.7 | 0.00 | 0 | 71.4 |
| JED-JLA | PSCex | 9 | -2364.8 | 4747.6 | 0.0 | 1.00 | 1 | 21.6 |
| | SCex | 9 | -2373.5 | 4765.1 | 17.5 | 1.00 | 0 | 35.4 |
| | SC | 6 | -2429.4 | 4870.8 | 123.1 | 0.97 | 0 | 326.0 |
| | PSC | 6 | -2470.7 | 4953.4 | 205.8 | 0.94 | 0 | 156.7 |
| | IM | 5 | -2630.5 | 5270.9 | 523.3 | 0.86 | 0 | 455.0 |
| | IMex | 8 | -2638.4 | 5292.8 | 545.2 | 0.85 | 0 | 228.8 |
| | SIex | 6 | -2880.9 | 5773.8 | 1026.1 | 0.72 | 0 | 411.4 |
| | SI | 3 | -2894.4 | 5794.8 | 1047.2 | 0.71 | 0 | 397.6 |
| | PAM | 6 | -2986.5 | 5984.9 | 1237.3 | 0.66 | 0 | 255.5 |
| | AM | 6 | -3088.9 | 6189.8 | 1442.2 | 0.61 | 0 | 125.0 |
| | AMex | 9 | -3441.9 | 6901.9 | 2154.3 | 0.41 | 0 | 59.9 |
| | PAMex | 9 | -4195.3 | 8408.6 | 3660.9 | 0.00 | 0 | 114.2 |
| JCA-JFR | PAMex | 9 | -271.6 | 561.2 | 7.8 | 0.93 | 0.02 | 16.2 |
| | SI | 3 | -273.7 | 553.3 | 0.0 | 1.00 | 0.93 | 327.5 |
| | IM | 5 | -274.9 | 559.8 | 6.4 | 0.94 | 0.04 | 364.6 |
| | AM | 6 | -275.1 | 562.1 | 8.8 | 0.92 | 0.01 | 436.0 |
| | AMex | 9 | -277.0 | 572.0 | 18.6 | 0.82 | 0.00 | 44.6 |
| | SIex | 6 | -280.3 | 572.5 | 19.2 | 0.82 | 0.00 | 317.9 |
| | IMex | 8 | -282.9 | 581.7 | 28.4 | 0.73 | 0.00 | 317.4 |
| | PSCex | 9 | -287.5 | 593.0 | 39.6 | 0.62 | 0.00 | 86.6 |
| | PAM | 6 | -296.8 | 605.7 | 52.3 | 0.50 | 0.00 | 405.4 |
| | SCex | 9 | -300.7 | 619.3 | 66.0 | 0.37 | 0.00 | 244.5 |
| | PSC | 6 | -301.8 | 615.6 | 62.2 | 0.41 | 0.00 | 371.6 |
| | SC | 6 | -323.3 | 658.6 | 105.3 | 0.00 | 0.00 | 157.0 |

K: The number of free parameters in the model

LogL: best maximum likelihood estimates over 20 independent runs

AIC: Akaike Information Criterion

$\Delta AIC_i$: Difference in AIC between model i and the best model

$W_{AIC}$: Akaike weight of model i compared to the best model

Theta: Theta parameter for the ancestral population before split ($\theta = 4N_{ref}\mu$), with $N_{ref}$ being the effective size of the ancestral population, and $\mu$ the per-site mutation rate per generation.

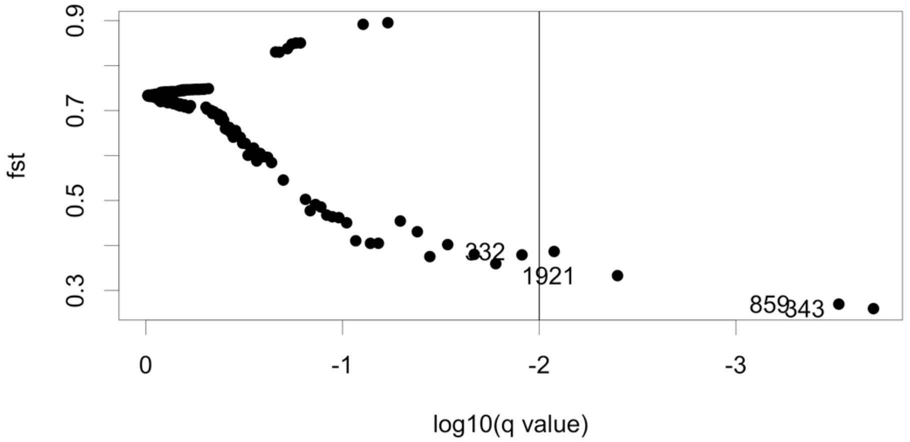


**Fig. S1.** Outlier analyses (BayeScan) based on 2,596 SNPs using 100 prior odds and 5000 pilot runs, followed by 100,000 iterations (5,000 samples, a thinning interval of 10, and a burn-in of 50,000).

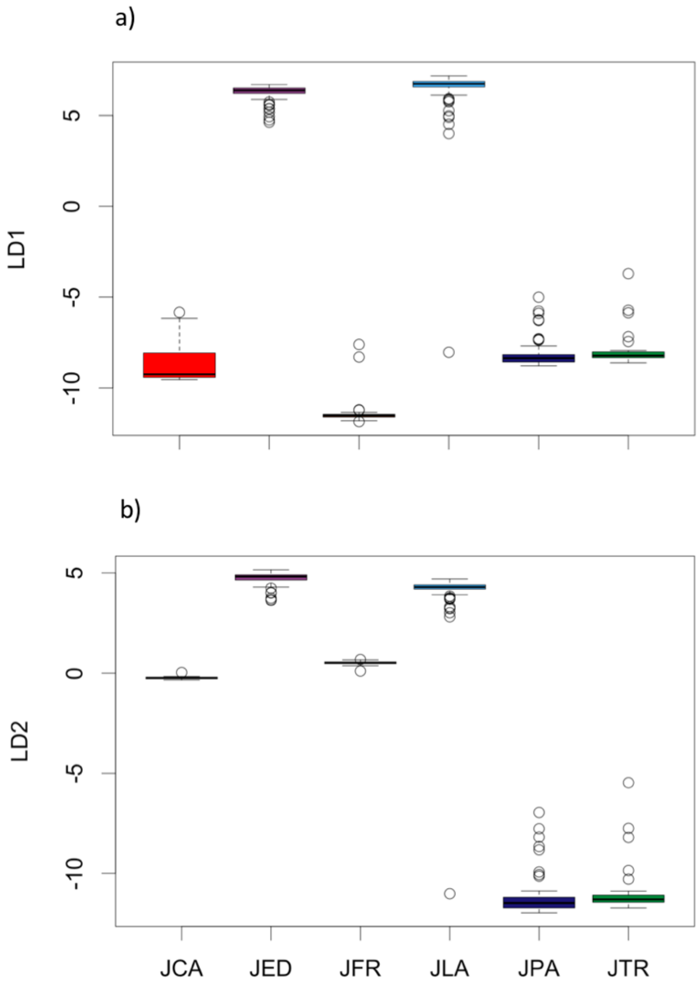

**Fig. S2.** Boxplots of individual coordinates along a) the first DAPC axis (LD1), which explained 29.9% of the variation and b) the second DAPC axis (LD2), which explained 22.4% of the variation. JCA: *J. caveorum*, JED: *J. edwardsii*, JFR: *J. frontalis*, JLA: *J. lalandii*, JPA: *J. paulensis*, JTR: *J. tristani*.

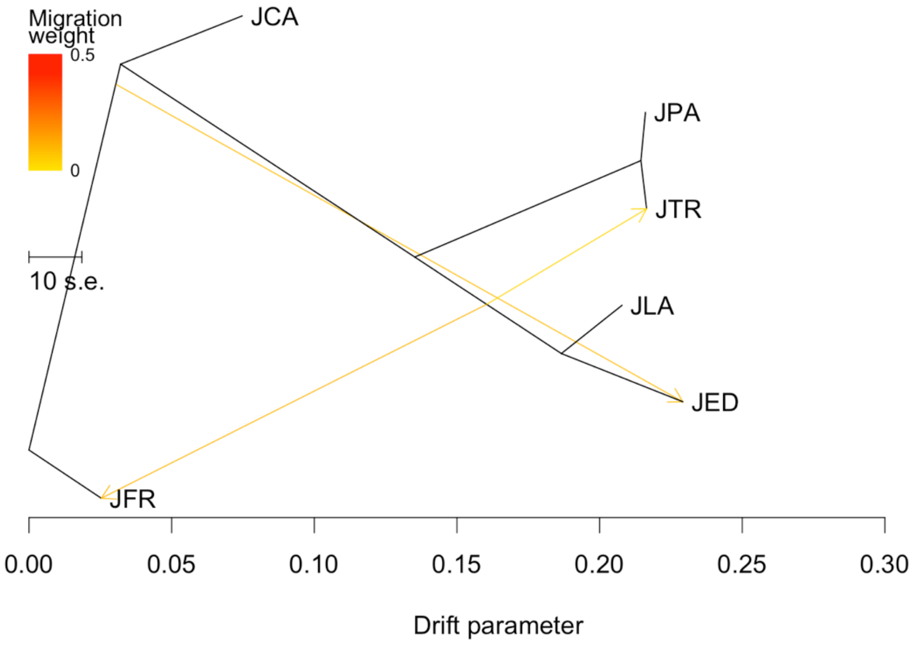


**Fig S3**. TreeMix results showing three ancestral admixture events.

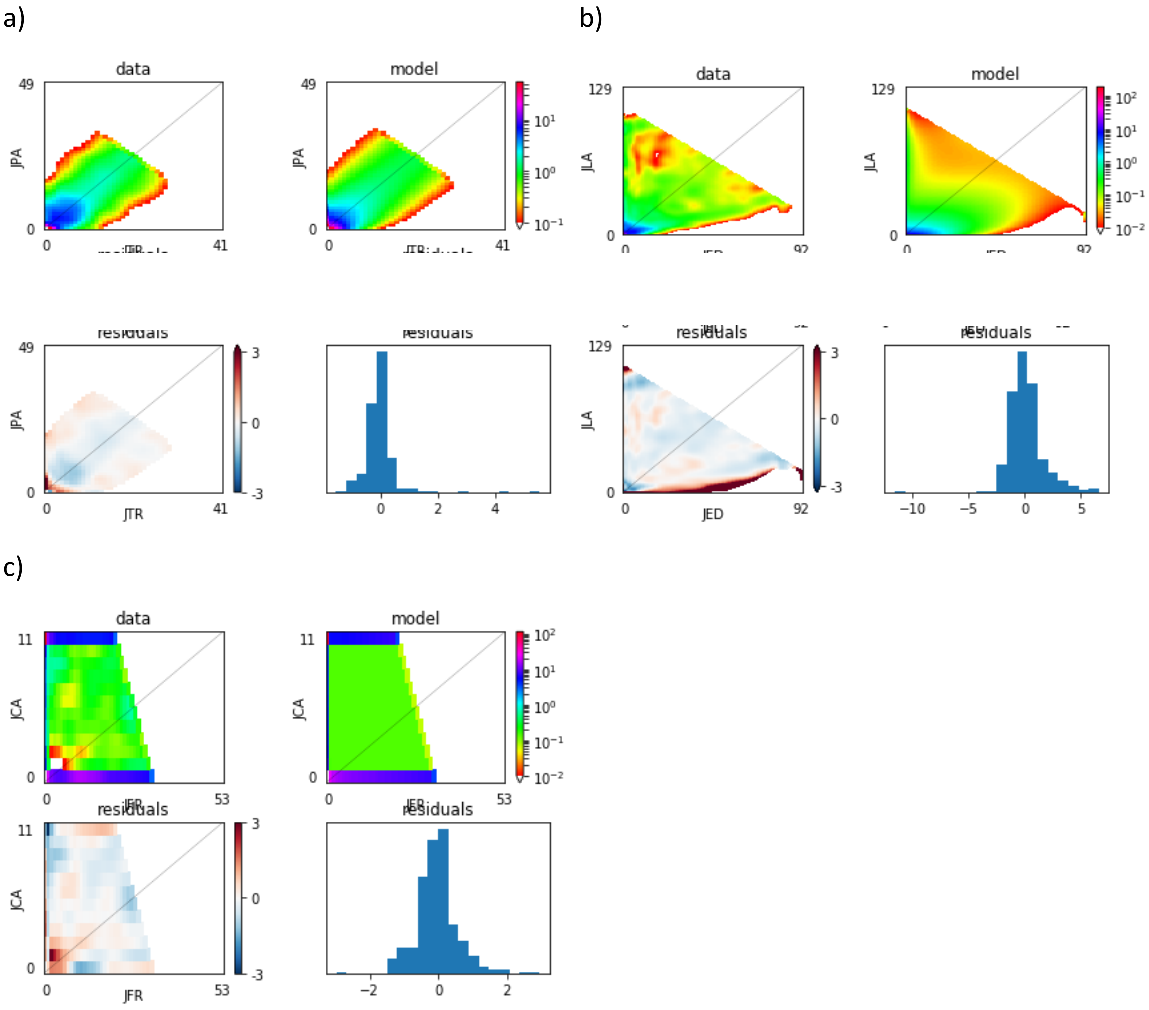


**Fig. S4.** Joint site frequency spectrum (JSFS) for a) *J. paulensis – J. tristani*, b) *J. lalandii – J. edwardsii*, and c) *J. caveorum – J. frontalis*. (When populations are isolated, each has its own segregating variants; this translates, in the joint SFS, into variants being restricted to the axes of the JSFS as it is the case of c)).

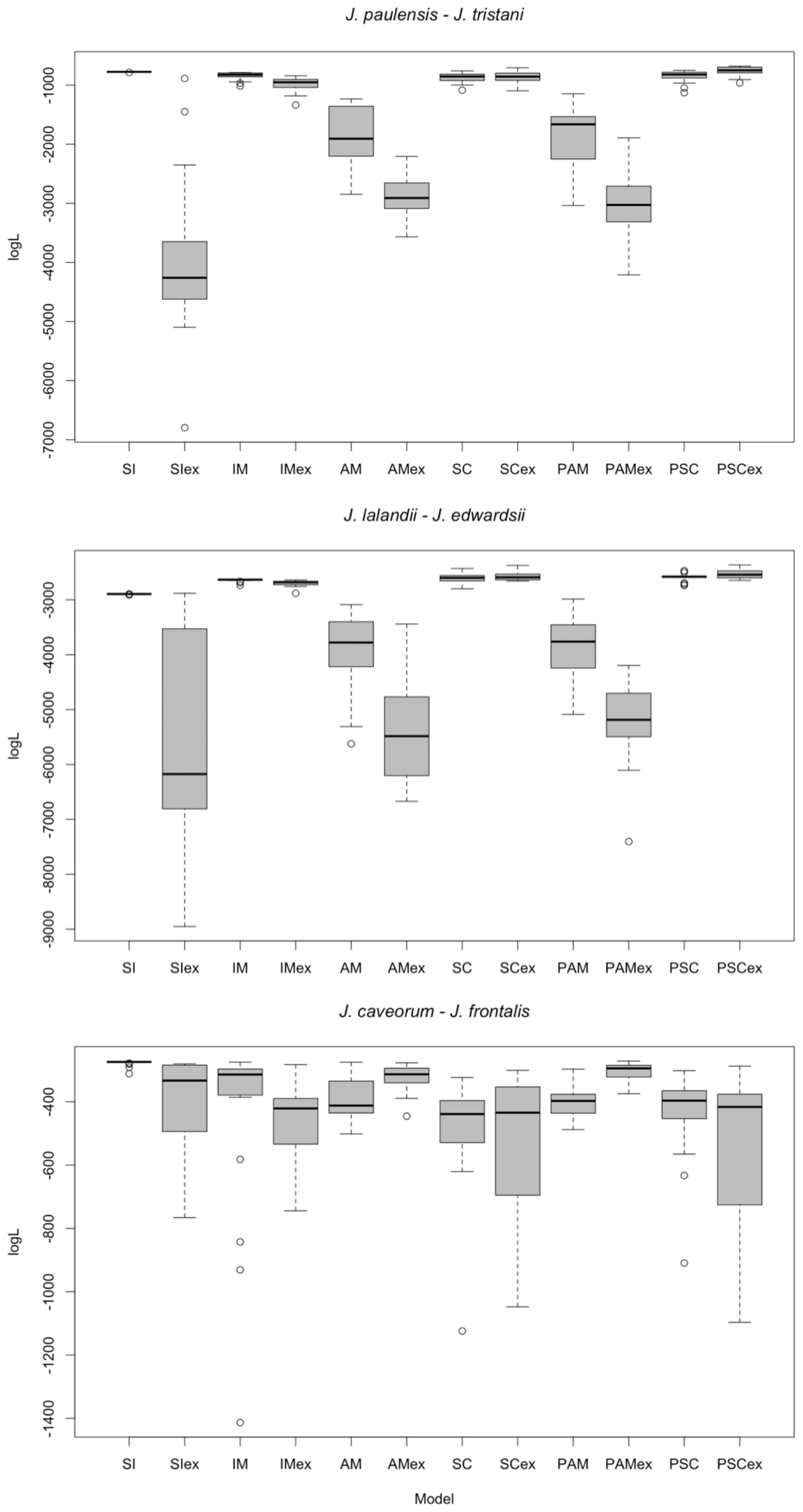


**Fig. S5.** LogL of 20 replicates for alternative models of divergence for each species pair. Strict Isolation (SI), Isolation-with-Migration (IM), Ancient Migration (AM), Secondary Contact (SC), Ancient Migration with two periods of ancient gene flow (PAM) and Secondary Contact with two periods of contact (PSC); and additional six models including population expansion (prefix 'ex').
